# Direct labeling of microtubule turnover reveals in-lattice repair and stabilization patterns in developing neurons

**DOI:** 10.64898/2026.01.11.698892

**Authors:** Ciarán Butler-Hallissey, Harrison M. York, Florence Pelletier, Jean-Marc Goaillard, Jérémie Gaillard, Manuel Théry, Pascal Verdier-Pinard, Christophe Leterrier

## Abstract

The microtubule cytoskeleton is the backbone of neuronal morphogenesis, driving the development of the dendrites and axon, and supporting trafficking to distant compartments. How neuronal microtubules are maintained and renewed in the face of this dynamic endeavor and long-term commitment remains unclear. Recently, in-lattice repair has emerged as an alternative mechanism of microtubule renewal that could allow for continuity of the existing network and for the emergence of cell polarity. Here, we directly assessed microtubule turnover using microinjection of labeled tubulin in cultured hippocampal neurons that exhibit well defined stages of development and polarization during the first 24 hours in culture. Sizeable tubulin integration was visible minutes after microinjection, indicating fast turnover of microtubules in developing neurons. Once it appeared, a longer neurite that would become the nascent axon showed decreased turnover, both for its acetylated and non-acetylated populations of microtubules. Combining microinjection with expansion microscopy allowed us to visualize tubulin integration sites along single microtubules, unambiguously demonstrating the existence of in-lattice integration along neurites. In thick neurites, we observed preferential integration within non-acetylated cortical microtubules, but in-lattice integration sites were also visible in the deeper core bundles of acetylated microtubules. Overall, our results link previous observations of microtubule stabilization patterns in developing neurons to their actual turnover. Mapping these patterns of turnover strengthens the notion that microtubules establish an organized network that participates in axon emergence and the establishment of neuronal polarity.

## Introduction

As a part of the cytoskeleton, microtubules are key players in cell organization, polarization, migration, division and intracellular transport (Akhmanova and Kapitein, 2022). They are formed by a head-to-tail polymerization of tubulin α/β heterodimers, which creates protofilaments that laterally interact to constitute the wall of a microtubule (Tilney et al., 1973). Consequently, the microtubule lattice is polarized with α-tubulins and β-tubulins exposed at the minus- and plus-end, respectively (Cleary and Hancock, 2021; McIntosh, 2025). Microtubules rapidly grow and collapse via a process known as dynamic instability, which occurs mostly at their plus-end in cells (Mitchison and Kirschner, 1984). To date, dynamic instability has been considered the main source of microtubule maintenance and renewal in cells (Gudimchuk and McIntosh, 2021).

The key roles of microtubules in morphogenesis, structural support and trafficking are even more critical in neurons displaying intricate shapes and striking polarization (Conde and Caceres, 2009; Kapitein and Hoogenraad, 2015; Penazzi et al., 2016). Microtubules form dense bundles within neurites (Iwanski and Kapitein, 2023), with their plus-end pointing uniformly away from the cell body in the axon, while the dendrites contain bundles of mixed plus-end and minus-end out orientations (Baas et al., 1988; Baas and Lin, 2011). Dendritic plus-end out microtubules are more dynamic, as demonstrated using plus-end interacting proteins (+TIP) tracking and laser ablation (Yau et al., 2016). Super-resolution microscopy recently showed that these plus-end out, dynamic microtubules were organized as a shell of tyrosinated microtubules in the vicinity of the plasma membrane, while a core of minus-end out, acetylated microtubules was assumed to be a more stable population (Tas et al., 2017; Katrukha et al., 2021).

Before the establishment of these patterns, microtubules are crucial during neuronal morphogenesis, with key roles in neurite growth and axon emergence (Beuningen and Hoogenraad, 2016; Higgs and Das, 2022). In the classical model of cultured hippocampal neuron development (Dotti et al., 1988; Banker, 2018), cells first develop peripheral lamellipodia (stage 1), then develop undifferentiated neurites after ∼24 hours (stage 2), before one of the neurites starts to grow faster and becomes the axon (stage 3). Interestingly, microtubules were thought to be uniformly oriented plus-end out in undifferentiated neurites and nascent dendrites (Baas et al., 1989), but more recent results challenged this view by showing that they contained bundles of mixed polarity from the earliest stages of neurite growth (Kollins et al., 2009; Iwanski et al., 2025).

In these studies, a lot of insights about microtubule stability patterns were inferred from drug resistance, post-translational modifications (acetylation), or +TIP dynamics, but these approaches do not directly assess microtubule turnover as a measure of their stability. This can be obtained using microinjection of labelled tubulin, which highlights newly formed microtubules if fixed and imaged shortly after injection (Soltys and Borisy, 1985; Schulze and Kirschner, 1986). Microinjection was performed in ganglionic neurons, selected for their large cell bodies, to distinguish newly assembled microtubules from pre-assembled ones within the axon, thereby demonstrating microtubule transport (Li and Black, 1996; Miller and Joshi, 1996; Yu et al., 1996; Slaughter et al., 1997). Later studies managed to microinject much smaller cortical neurons of Syrian hamsters, imaging dynamics of microtubules at axonal branches and growth cones hours later (Dent et al., 1999; Dent and Kalil, 2001). These studies did not address microtubule turnover due to their slow time course and owing to the difficulty in visualizing single microtubules in densely packed neurites, which required perturbative preparation methods such as microtubule “fraying” induced by high salt concentration before fixation (Li and Black, 1996).

Directly assessing neuronal microtubule turnover is made necessary by recent results about the fundamental mechanisms of microtubule turnover. Microtubules can not only be created de novo (nucleation) or renewed by shrinkage and growth from their plus-end (end-renewal) but can also be renewed via the replacement of tubulin dimers within the assembled lattice that results from damage (dimer loss) followed by repair (in-lattice insertion, Théry and Blanchoin, 2021; Motta et al., 2023). These damages can be induced by mechanical stress (Schaedel et al., 2015), activity of severing enzymes (Vemu et al., 2018), or motor proteins travel (Triclin et al., 2021; Budaitis et al., 2022; Andreu-Carbó et al., 2024), but also happen spontaneously (Schaedel et al., 2019). We recently demonstrated that microinjection of labelled tubulin shortly before fixation was a method of choice to visualize in-lattice insertion (Gazzola et al., 2023), resulting in better specificity and sensitivity than the photoconversion approach previously used (Aumeier et al., 2016).

The higher stability of neuronal microtubules (Baas et al., 2016) and the importance of preserving an intact network for transporting cargoes along neuronal extensions make in-lattice renewal a compelling mechanism for microtubule turnover in neurons, allowing for “in-place” replacement of microtubules akin to the eternal ship of Theseus (Guedes-Dias and Holzbaur, 2019) and for cellular polarization via the specification of highly-repaired microtubule arrays (Théry and Blanchoin, 2021; Andreu-Carbó et al., 2022). We thus decided to directly investigate microtubule turnover and assess the presence of in-lattice integration by performing microinjection of labelled tubulin in developing neurons, using the well-characterized model of dissociated hippocampal cultures (Banker, 2018). We focused on early development after one day in vitro (1 DIV) and microinjected biotinylated tubulin into stage 1 (lamellipodia), stage 2 (undifferentiated neurites) and stage 3 (specified, longer axon) neurons (Dotti et al., 1988). Fixing cells within 25 minutes after microinjection and quantifying the turnover in distinct stages and in different compartments revealed that nascent axons show a lower microtubule turnover compared to minor neurites. We next combined microinjection of tubulin with expansion microscopy (ExM) to resolve the distinct turnover rate within the membrane-apposed shell microtubules and the acetylated core bundles, and to unambiguously reveal the presence of in-lattice insertion besides end-renewal as a source of microtubule turnover in neurons.

## Results

### Microinjection of biotinylated tubulin in immature neurons reveals fast integration in microtubules via plus-end growth and in-lattice renewal

To directly reveal events of tubulin integration in assembled microtubules, we microinjected labeled tubulin into the cell body of rat hippocampal neurons after 22-27 hours in culture (1 DIV). To overcome the small cytoplasmic volumes when microinjecting these neurons compared to previously used ganglionic neurons (Li and Black, 1996), we fabricated nano-pipettes with a tip outer diameter of less than 0.3 µm (see Methods and Fig. S1A). A workflow for microinjection of biotinylated tubulin with dextran-FITC was implemented to monitor the introduction of tubulin into the neurons (see Methods, Movie S1). Out of 124 injected neurons, the majority (65%) were fixed between 0 and 10 minutes after microinjection (binned integration durations, Fig. S1B-C), to detect fast integration of tubulin into existing microtubules. After fixation, neurons were immunostained for microtubules (α-tubulin, αTub) and for acetylated microtubules (acetylated α-tubulin, acTub), and for the microinjected and integrated tubulin (intTub).

Super-resolution spinning-disk confocal microscopy of these microinjected and stained immature neurons revealed robust tubulin integration along microtubules within minutes of microinjection (Fig. 1A). In areas such as neurite tips or cell body where individual microtubules were resolvable, we observe bright stretches of integrated tubulin at the tip of microtubules, whereas fainter punctate integration spots were present along the lattice (Fig. 1A-B). These patterns are similar to the ones observed in non-neuronal cells (Gazzola et al., 2023) and are likely corresponding to plus-ends growth and in-lattice insertion events, respectively. At longer time points after microinjection, the density of tubulin integration made it difficult to distinguish individual integration events at this imaging resolution (Fig. 1C).

**Figure 1.**
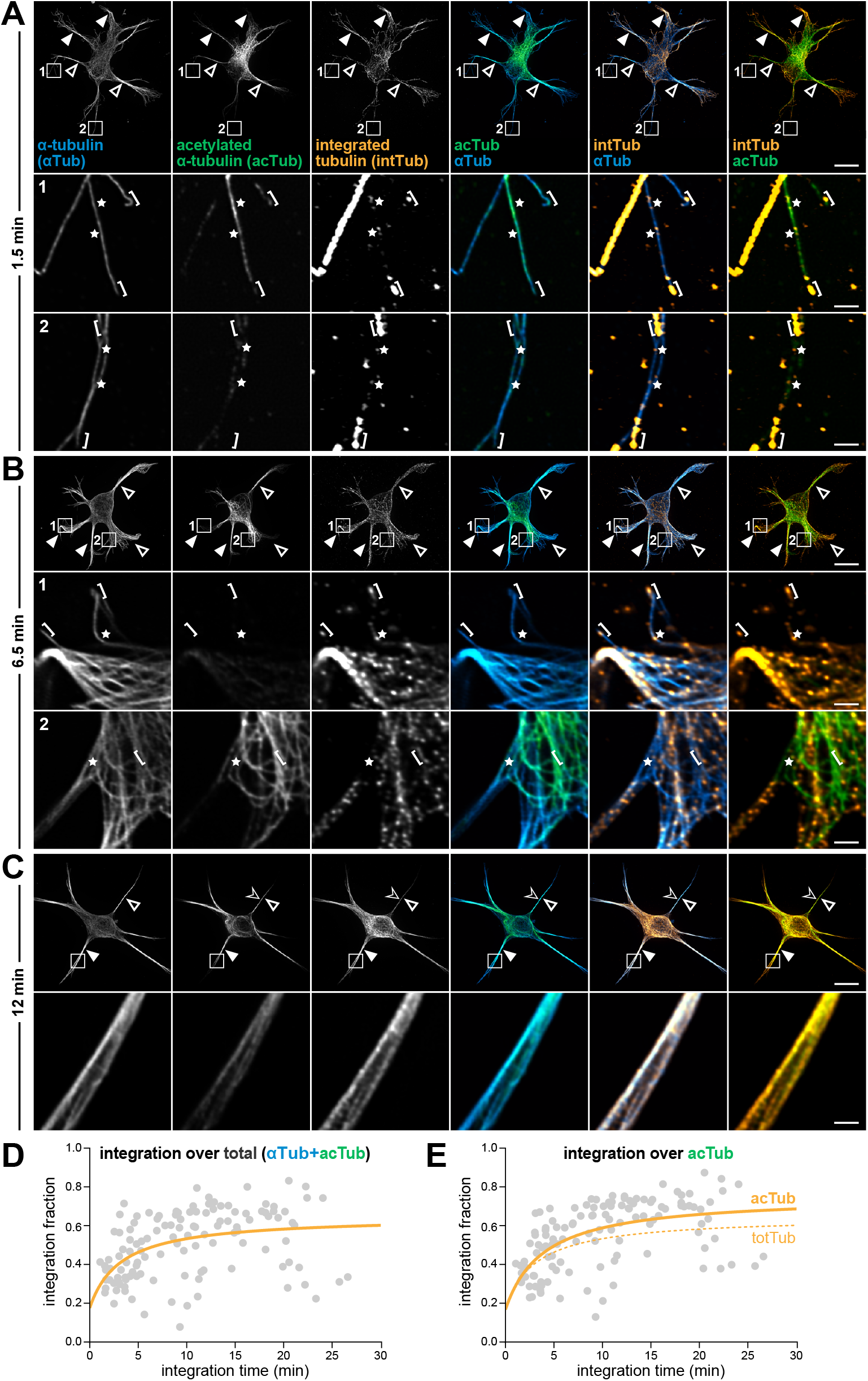
Microinjection of biotinylated tubulin in immature neurons reveals fast integration in microtubules via plus-end growth and in-lattice renewal. **A**. Rat hippocampal neuron at 1 DIV fixed after microinjection with biotinylated tubulin, fixed after a tubulin integration time of 1.5 minutes, and stained for α-tubulin (αTub, blue on overlays), acetylated α-tubulin (acTub, green on overlays) and integrated tubulin (intTub, orange on overlays). Images are deconvolved super-resolution spinning disk maximum intensity projections. On whole neuron images, solid white arrows show areas with high integrated tubulin signal, while hollow white arrows show areas of lower integrated tubulin signal. Zooms show the area boxed on the whole image as a single Z slice, with white stars pointing putative in-lattice tubulin integrations, and white brackets indicating putative growing plus-ends. Scale bars: 10 µm (whole neuron images), 1 µm (zooms). **B**. 1 DIV neuron fixed after a tubulin integration time of 6.5 minutes, similar to A. **C**. 1 DIV neuron fixed after a tubulin integration time of 12 minutes, similar to A. **D**. Graph of the integration fraction of microinjected tubulin for the whole injected neuron population as a function of integration time. Quantification of all microinjected neurons showing the integration fraction over the total tubulin positive pixels (αTub+acTub). Orange curve is a fit using nonlinear regression (Padé (1,1) approximant equation). **E**. Graph of the integration fraction as a function of integration time for microinjected tubulin for the whole injected neuron population, over the acetylated α-tubulin positive pixels. Orange curve is a fit similar to D, with the dashed curve being the same fit as D for comparison. N = 124 neurons for D and E.

We next quantified the rate of integration of microinjected tubulin over time. We calculated the fraction of pixels positive for integrated tubulin that overlap with the total tubulin mask (defined as the sum of the α-tubulin and acetylated α-tubulin segmented masks, Fig. S2A) or the acetylated α-tubulin mask over the whole neuron (Fig. 1D-E and Fig. S2B). This integration fraction was plotted against the integration time, defined as the time between microinjection and fixation (Fig. S1B and S2B, output 1), to assess microtubule turnover in these cells. In less than 5 min, the integration fraction reached 40%, which is consistent with a fast tubulin integration rate, and plateaued at around 60% within 10 min (Fig. 1B). The whole-neuron integration curve was similar when integration was assessed over the total microtubule or the acetylated microtubule network, although the two plots were better fit by distinct curves (Fig. 1C).

### Whole-cell microtubule turnover shows similar kinetics across stages of neuronal development

We next took advantage of the presence of distinct early developmental stages in our microinjected neuron population at 1 DIV and tested if microtubule turnover could vary between these stages. Stages were defined as stage 1, stage 2 symmetric (2s), stage 2 asymmetric (2a) and stage 3 based the established developmental timeline of cultured hippocampal neurons (Dotti et al., 1988), with an added distinction between symmetrical and asymmetrical stage 2 neurons (see Methods and Fig. S3A). Out of 124 microinjected neurons, only 4 were classified as stage 1, 59 as stage 2s, 39 at stage 2a, and 22 at stage 3 (Fig. S3B). At all stages, we could identify examples of plus-end growth as bright stretch of integrated tubulin at the end of a microtubule, and in-lattice insertions as fainter spots within a continuous microtubule (Fig. 2A-D).

**Figure 2.**
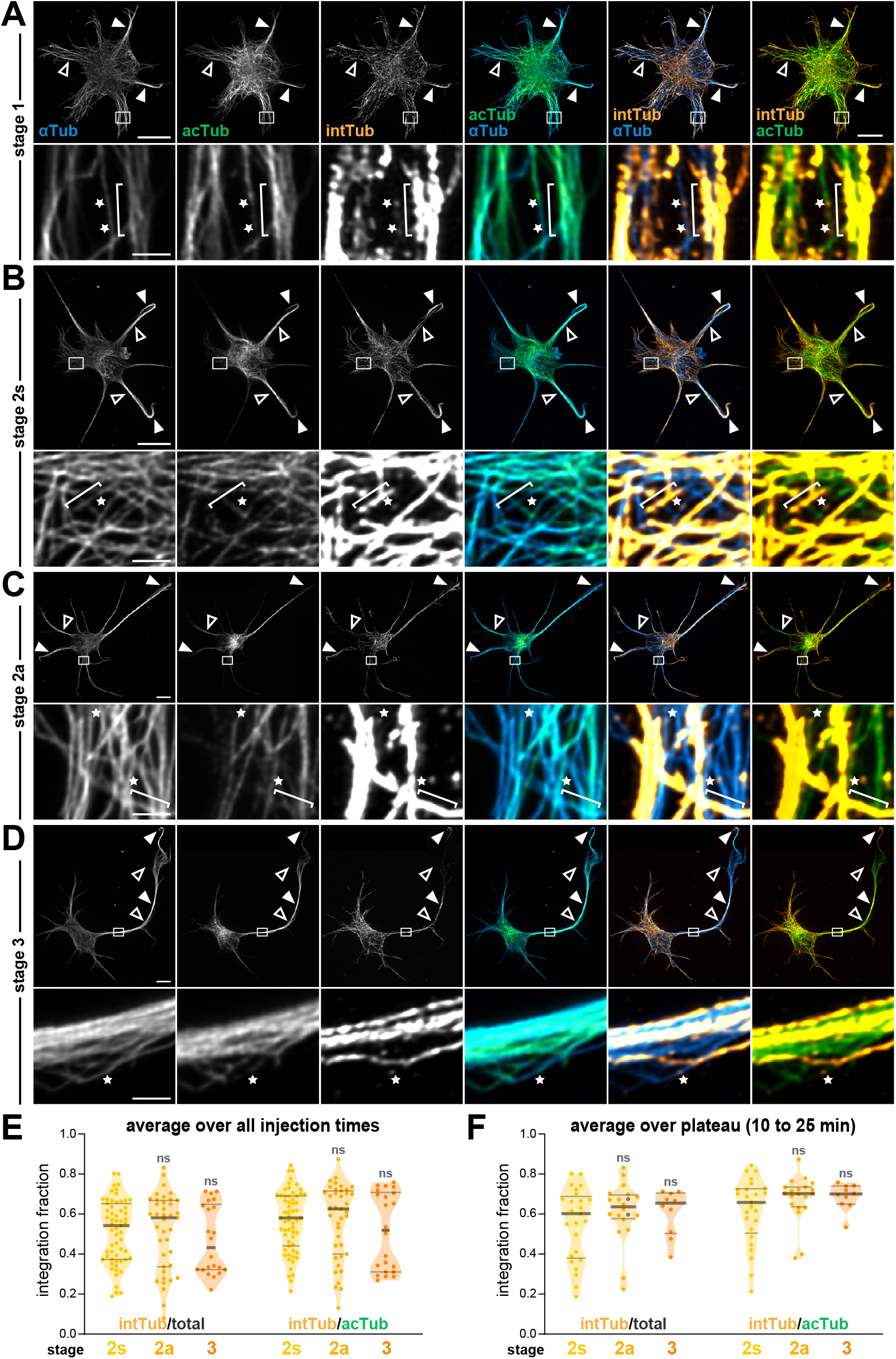
Whole-cell microtubule turnover shows similar kinetics across stages of neuronal development. **A**. Deconvolved super-resolution spinning disk maximum intensity projections of a 1 DIV neuron at developmental stage 1 (see Methods and Fig. S3A), microinjected and fixed after a tubulin integration time of 3 to 4 minutes, stained for α-tubulin (αTub, blue on overlays), acetylated α-tubulin (acTub, green on overlays) and integrated tubulin (intTub, orange on overlays). Solid white arrows show areas with a high integrated tubulin signal, while hollow white arrows show areas with a lower integrated tubulin signal. Zooms show the area boxed on the whole image as a single Z slice, white stars pointing putative in-lattice tubulin integrations, and white brackets indicating putative growing plus-ends. Scale bars: 10 µm (whole neuron images), 1 µm (zooms). **B**. 1 DIV neuron at developmental stage 2s, fixed 3 to 4 minutes after injection, similar to A. **C**. 1 DIV neuron at developmental stage 2a, fixed 3 to 4 minutes after injection, similar to A. **D**. 1 DIV neuron at developmental stage 3, fixed 3 to 4 minutes after injection, similar to A. **E**. Graph of the integration fraction of microinjected tubulin for the whole injected neuron population, averaged over all integration times for developmental stages 2s, 2a and 3. The integration fraction is calculated over the total tubulin positive pixels (left) or over the acetylated α-tubulin positive pixels (right). **F**. Graph of the integration fraction of microinjected tubulin for the whole injected neuron population averaged over integration times between 10 and 25 minutes, similar to E. For E and F, violin plots show the median (thick horizontal segment) and upper 75th and lower 25th interquartile ranges (thin horizontal segments). Statistical analysis was performed using the Kruskal–Wallis nonparametric comparing across developmental stages for total tubulin or acetylated tubulin, with significance testing between a developmental stage and the one just to its left (previous stage). N=22-59 neurons per stage, 12-23 neurons per stage (F).

Analyzing the kinetics of tubulin integration at distinct stages showed that the average integration fraction (across injection times) was not different for stages 2s, 2a and 3, although a slight drop was seen at stage 3 (Fig. 2E). The integration fraction plateaued between 10 and 25 minutes, and the level of this plateau was not different between stage 2s, stage 2a, and stage 3 (Fig. 2F). At this level, we also found no difference between the integration rate over the total microtubule network compared to the acetylated microtubule one (Fig. S4). We could detect integration at the tip of the longest neurites less than five minutes post-microinjection, suggesting that diffusion of microinjected tubulin was not a limiting factor for the integration in regions distal from the cell body (Fig. 2A-D).

### Microtubules gradually accumulate newly incorporated tubulin in neurites during development

The fast and heterogeneous microtubule turnover at every developmental stage prompted us to perform a quantitative analysis of tubulin integration within neuronal compartments. To do this, we set up a two-level analysis that allowed to measure the enrichment of acetylated tubulin or integrated tubulin in each compartment. At the first level, we segmented the cell body and neurites of each neuron (Fig.3A and S2C-D) and the intensities of the pixels positive for α-tubulin, acetylated α-tubulin, or integrated tubulin were summed over each compartment, then expressed as a percentage of the total signal within the neuron (integrated density for all compartments). This distribution percentage for α-tubulin, acetylated tubulin and integrated tubulin measures the relative distribution of the label in the neurites and in the cell body (Fig. S2E, output 2). At the second level, in each compartment we calculated the distribution ratio for acetylated α-tubulin and integrated tubulin by dividing their distribution percentages by the α-tubulin distribution percentage, providing the relative enrichment of tubulin acetylation and tubulin integration in the cell body and neurites (Fig. S2E, output 3). We further stratified these distribution percentages and ratios for each compartment across stages before (2s), during (2a) and after (3) axon emergence (Fig. 3B-F).

**Figure 3.**
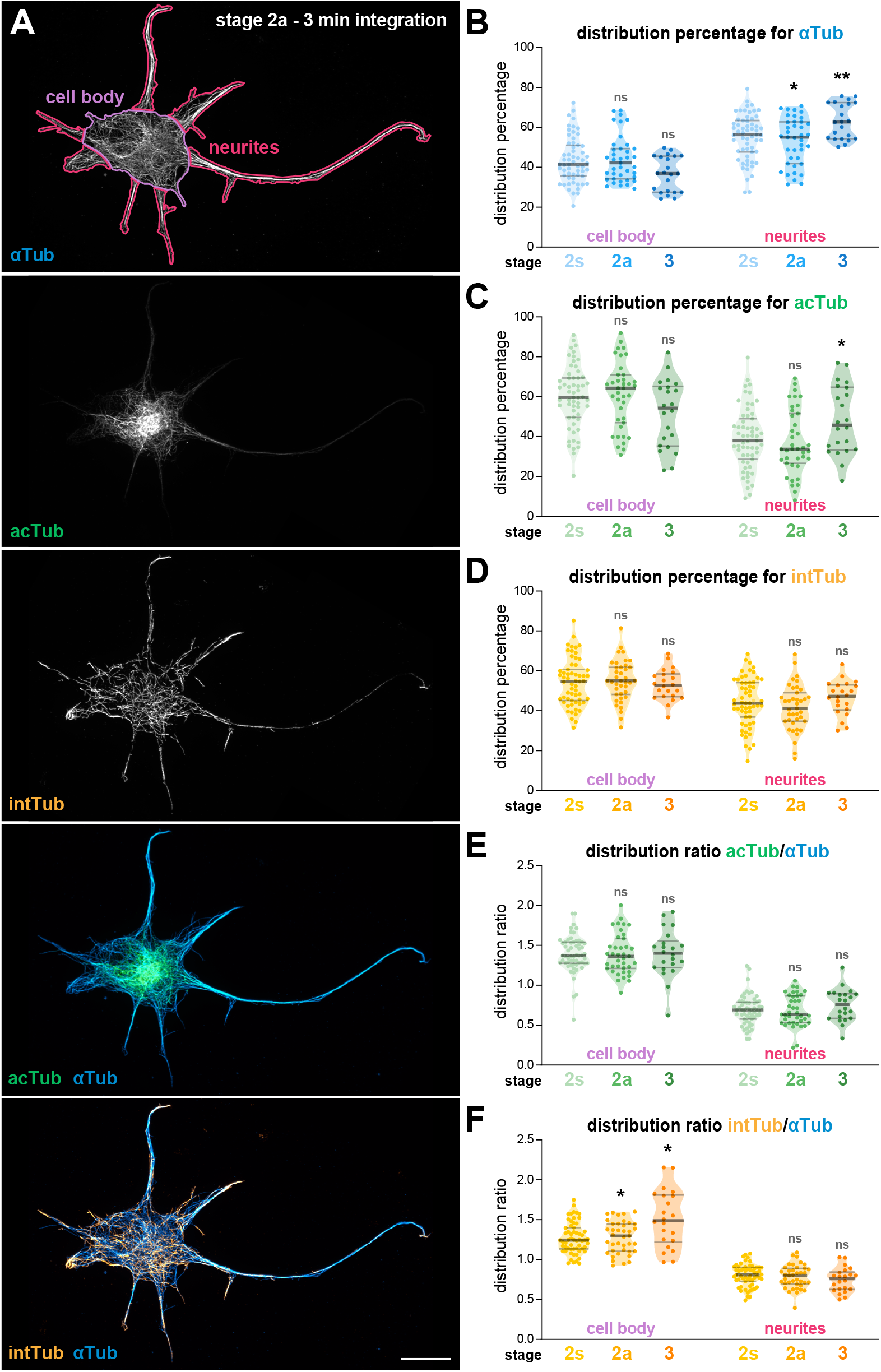
Microtubules gradually accumulate newly incorporated tubulin in neurites during development. **A**. Deconvolved super-resolution spinning disk maximum intensity projections of a 1 DIV neuron at stage 2a, microinjected and fixed after 3 minutes of tubulin integration, stained for α-tubulin (αTub, blue on overlays), acetylated α-tubulin (acTub, green on overlays) and integrated tubulin (intTub, orange on overlays). Magenta and purple compartments on the αTub image highlight the segmentation of the cell body and neurites, respectively. Scale bar, 10 µm. **B**. Graph of the distribution percentages for α-tubulin (αTub) measuring its relative distribution in the cell body and neurites by developmental stage. All neurites for a given neuron were considered as one compartment. **C**. Graph of the distribution percentages for acetylated α-tubulin (acTub) measuring its relative distribution in the cell body and neurites by developmental stage, similar to B. **D**. Graph of the distribution percentages for integrated tubulin (intTub) measuring its relative distribution in the cell body and neurites by developmental stage, similar to B. **E**. Graph of the distribution ratios for acetylated α-tubulin (acTub) measuring its enrichment in the cell body and neurites compared to α-tubulin (αTub) by developmental stage. All neurites for a given neuron were considered as one compartment. **F**. Graph of the distribution ratios for acetylated α-tubulin (acTub) measuring its enrichment in the cell body and neurites compared to α-tubulin (αTub) by developmental stage, similar to E. For B to F, violin plots show the median (thick horizontal segment) and upper 75th and lower 25th interquartile ranges (thin horizontal segments). Statistical analysis was performed using the Kruskal–Wallis nonparametric test followed by Dunn’s multiple comparisons test, with significance testing between a developmental stage and the one just to its left (previous stage). N = 22-59 (cell bodies), 116-274 (neurites).

At the level of single label distribution, the majority of microtubule (α-tubulin distribution percentage) were found in neurites, with a significant increase in stage 3 neurons reflecting neurite growth (Fig. 3B, median αTub distribution percentage in neurites is 56% for stage 2s, 55% for stage 2a, and 63% for stage 3). The major proportion of acetylated microtubules was found in the cell body throughout the stages, but the distribution also increased toward neurites along maturation (Fig. 3C, acTub distribution percentage in neurites: 38% for stage 2s, 34% for stage 2a, and 46% for stage 3). While the cell body was the site of microinjection, on average, the integrated tubulin distribution ratio for the cell body was only ∼10% higher than for the neurites. There was no significant difference in integrated tubulin distribution ratios between neurons of different stages, either in the cell body or the neurites taken as a whole (Fig. 3D, intTub distribution percentage in neurites: 44%, 41% and 47% for stage 2s, 2a and 3, respectively).

We then derived the distribution ratios to measure how the enrichment of acetylated or integrated tubulin varied between the cell body and neurites as a whole during early neuronal development. This enrichment was constant across stages in the cell body and neurites for acetylated tubulin (Fig. 3E, distribution ratios in the cell body: 1.37, 1.36, 1.40; in the neurites: 0.69, 0.63, 0.76 for stage 2s, 2a and 3, respectively). This shows that at this broad subcellular level, no shift in relative acetylation levels occurs between the cell body and neurites. For integrated tubulin, the cell body enrichment increased from stage 2s and 2a (1.25 and 1.30, respectively) to stage 3 (1.49, Fig. 3F). This suggests that axon emergence is associated with higher tubulin turnover is cell body, whereas the global turnover in neurites is stable.

### Microtubule turnover in the longest neurite decreases upon axon emergence

Because the loss of neuron symmetry is due to the emergence of a permanently longer neurite differentiating into the axon between stage 2 and stage 3 (Fig. 4A-B and Fig. S3A), we refined our previous analysis to separate the longest neurite (nascent axon) from the shorter neurites (minor neurites). We measured neurite length from segmented images of 1 DIV neurons using a geodesic map (see Methods and Fig. S2F, output 4). The average length of neurites did not significantly change between stage 2a, 2s and 3, because most neurites are minor ones; but the length of the longest neurite increased steadily from stage 2s to 3 (Fig. S5A). We then plotted the distribution percentages of α-tubulin, acetylated tubulin and integrated tubulin in the minor neurites vs the longest neurite at different stages. As expected, the percentage in the longest neurite rose between stage 2s, 2a and 3 while the percentage in the minor neurites decreased (Fig. S5B), reflecting the selective growth of the nascent axon at stage 2a and 3.

**Figure 4.**
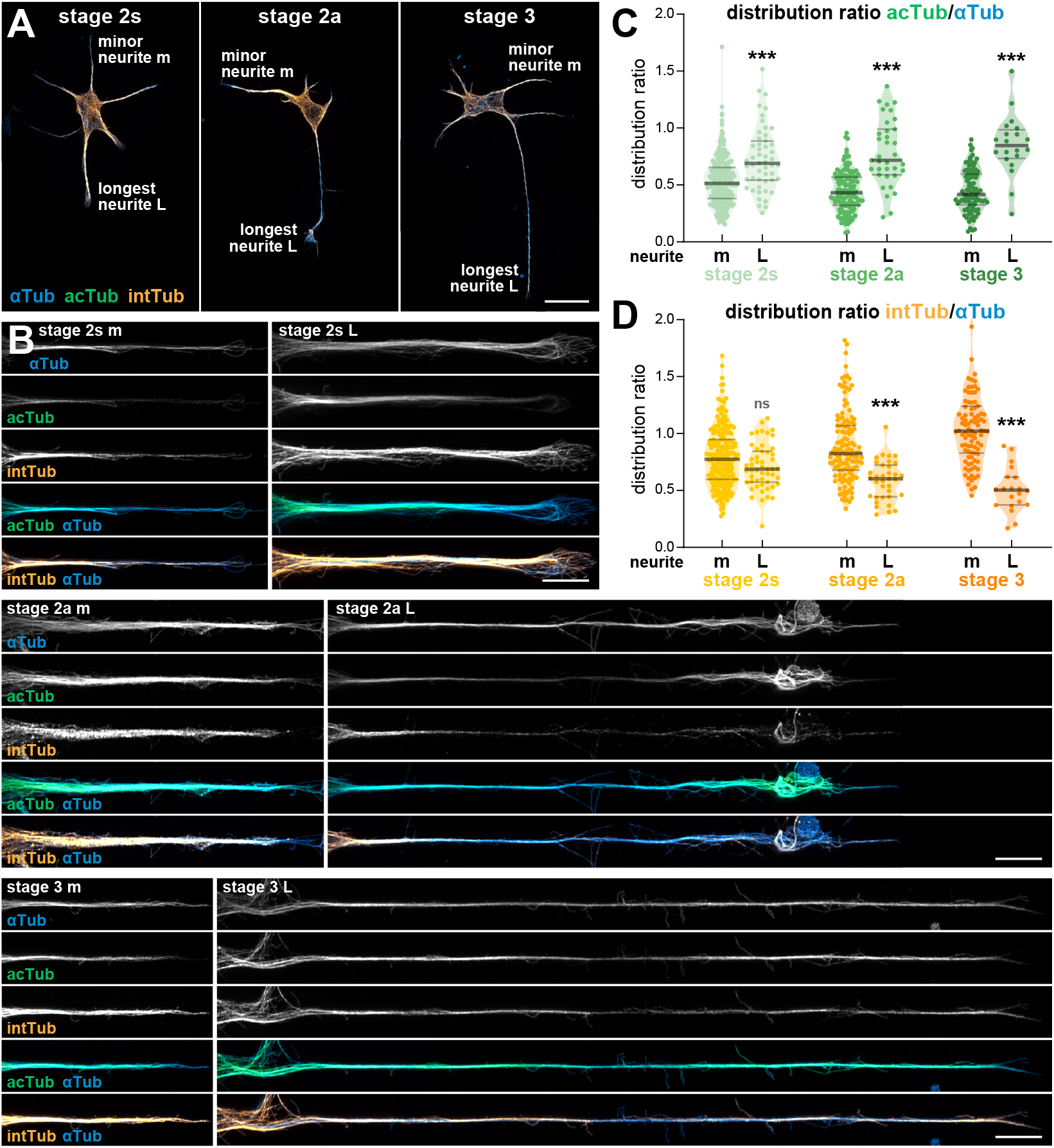
Microtubule acetylation increases while tubulin turnover decreases in the longest neurite upon axon emergence. **A**. Deconvolved super-resolution spinning disk maximum intensity projections of 1 DIV neurons at stage 2s (left), 2a (center) and 3 (right), microinjected and fixed after 11 to 12 minutes of tubulin integration, stained for α-tubulin (αTub, blue on overlays), acetylated α-tubulin (acTub, green on overlays) and integrated tubulin (intTub, orange on overlays). One minor neurite (m) and the longest neurite (L) are indicated. Scale bar, 20 µm. **B**. Minor (m) and longest (L) neurites indicated in A for each neuron at stage 2s; 2a and 3, zoomed and straightened. Scale bars, 5 µm. **C**. Graph of the distribution ratios for acetylated α-tubulin (acTub) measuring its enrichment in the minor neurites (m) and longest neurite (L) compared to α-tubulin (αTub) by developmental stage. Each neurite was considered individually, and branched neurites were not considered in the analysis (74 over a total of 648 neurites across developmental stages). **D**. Graph of the distribution ratios for acetylated α-tubulin (acTub) measuring its enrichment in the minor neurites (m) and longest neurite (L) compared to α-tubulin (αTub) by developmental stage, similar to C. For C and D, violin plots show the median (thick horizontal segment) and upper 75th and lower 25th interquartile ranges (thin horizontal segments). Statistical analysis was performed using the Kruskal–Wallis nonparametric test followed by Dunn’s multiple comparisons test, with significance testing between the longest neurites and the minor neurites at the same stage (just to its left). Stage 2s: minor neurites N = 223, longest neurites N = 51; Stage 2a: minor neurites N = 136, longest neurites N = 35; Stage 3: minor neurites N = 96, longest neurites N = 20.

The distribution ratios then allowed us to compare the relative enrichment of tubulin acetylation and integration in the longest neurite to the minor ones (Fig. 4C). We found that tubulin acetylation is selectively enriched in the longest neurites as soon as the symmetrical 2s stage, where differences in length are minimal and a shorter neurite may still emerge as the axon. This difference in acetylated tubulin enrichment between minor and longest neurites increased at stages 2a and 3 (acetylated tubulin distribution ratio minor/longest neurites: 0.51/0.69 at stage 2s, 0.43/0.71 at stage 2a, 0.41/0.84 at stage 3, Fig. 4C), in line with the previously observed higher acetylation of microtubules in the nascent axon, which was interpreted as a sign of microtubule stabilization (Witte et al., 2008).

Here, we could measure the microtubule turnover as the integration of the microinjected tubulin: the distribution ratio of integrated tubulin was slightly lower, but not significantly different in the longest neurite at stage 2s (0.77/0.69 for longest and minor neurites). This difference increased and became significant at stage 2a and 3 (0.82/0.60 at stage 2a, 1.02/0.50 at stage 3, Fig. 4D), directly demonstrating that the nascent axon exhibits a lower tubulin turnover and higher microtubule stability. Another way to demonstrate the higher acetylation and lower turnover of longer neurites is plot the acetylated or integrated tubulin distribution ratio as a function of the neurite length (Fig. S5C-D) for the whole dataset of 124 neurons: this reveals a positive correlation between tubulin acetylation and length (Spearman’s ρ value of 0.49), but a negative correlation between tubulin integration and neurite length (ρ value of -0.30, Fig. S5C-D). Altogether, our analyses suggests that a decrease in the tubulin turnover in the microtubules of the longest neurite correlates with the emergence of the axon.

### Expansion microscopy on microinjected neurons helps visualize single microtubules and tubulin integration in dense regions

Next, we wanted to go further to discriminate how tubulin is integrated in the existing microtubule network. Using super-resolution confocal microscopy, we saw bright stretches at microtubule extremities and small spots of integrated tubulin along microtubules in areas with low to moderate microtubule density, suggesting that plus-ends growth and in-lattice incorporation are present in neurons (see Fig. 1A-C and Fig. 2A-D). To distinguish those within the dense microtubule bundles along neurites, we subjected neurons microinjected with biotinylated tubulin to ultrastructure expansion microscopy (U-ExM, Gambarotto et al., 2018). Microinjected neurons were fixed, embedded in and expanded in a custom silicone well, stained for α-tubulin, acetylated tubulin and integrated tubulin, then relocated (see Methods).

We observed that microinjected tubulin was rapidly integrated in all neuronal compartments, with heterogenous integration patterns (Fig. 5, showing a stage 2s neuron). Thanks to U-ExM, we were able to distinguish putative plus-end growth and in-lattice integration not only in sparse microtubule areas (Fig. 5B-C), but also in denser regions within neurites and growth cones (Fig. 5D-E). Stretches of integration, likely representing plus-end growth at the extremity of a microtubule, appeared as a pattern of closely-apposed dots after expansion. We could also see a sparser distribution of tubulin integration dots along individualized segments of microtubules, most likely corresponding to in-lattice integration into existing microtubules (Fig. 5). Thus, U-ExM confirmed our super-resolved confocal images, extending the identification of in-lattice integration events to the microtubules along immature neurites.

**Figure 5.**
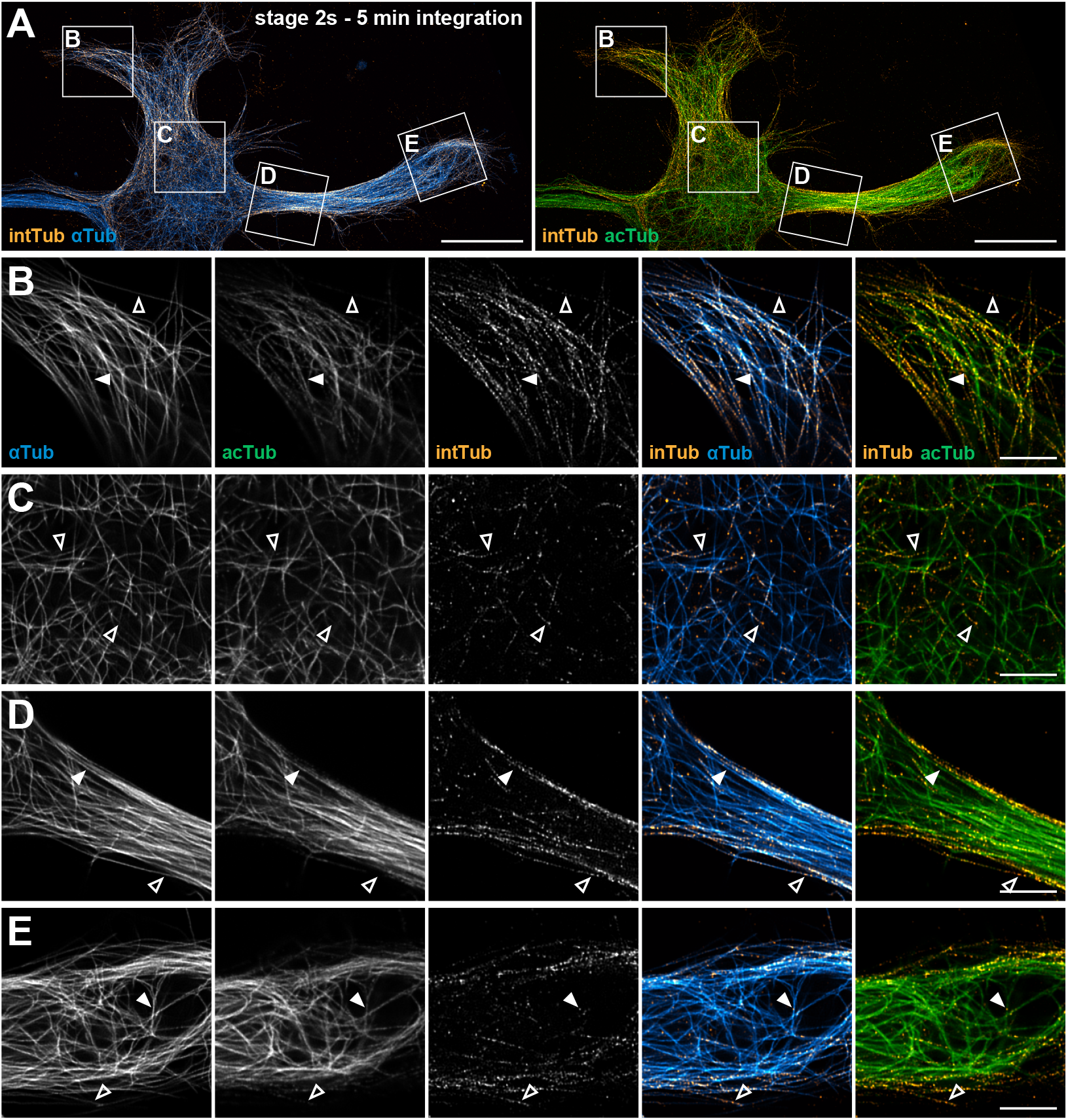
Expansion microscopy on microinjected neurons helps visualize single microtubules and tubulin integration in dense regions. **A**. Deconvolved spinning disk maximum intensity projections of a 1 DIV neuron at stage 2s, microinjected and fixed after 5 minutes of tubulin integration, processed for U-ExM and stained for α-tubulin (αTub, blue on overlays), acetylated α-tubulin (acTub, green on overlays) and integrated tubulin (intTub, orange on overlays). Scale bars, 10 µm after correction for the expansion factor. **B**. Zoom on a portion of a lamellipodia corresponding to a single Z-slice for the area boxed in A. Solid white arrowheads point to putative in-lattice tubulin integrations in acetylated microtubules, while hollow white arrowheads point to putative in-lattice tubulin integrations in non-acetylated or low-acetylated microtubules. **C**. Zoom on a portion of the cell body corresponding to a single Z-slice for the area boxed in A, similar to B. **D**. Zoom on a proximal portion a thick neurite corresponding to a single Z-slice for the area boxed in A, similar to B. **E**. Zoom on a proximal portion a thick neurite corresponding to a single Z-slice for the area boxed in A, similar to B. Scale bars, 2 µm after correction for the expansion factor.

Tubulin integration sites were present on both acetylated and non-acetylated microtubules, but in expanded proximal sections of neurites, our images showed a clear difference in the density of tubulin integration sites between the non-acetylated and acetylated microtubules located at the periphery and center of neurites, respectively (Fig.5D, Tas et al., 2017; Iwanski et al., 2025). The peripheral microtubules had a high density of integration sites and putative plus-end growth patterns, whereas few were present in central microtubules, which suggests a significantly higher turnover of microtubules at the periphery than at the center.

### Microtubules show more turnover at the peripheral shell of neurites, and less within the neurite core

To get a more precise view of this difference in turnover between peripheral and core microtubules in neurites, we performed 3D reconstructions of super-resolution confocal images of microinjected, expanded neurons (Movie S2). Cross-sections of large neurites in 1 DIV neurons confirmed that most of the integrated tubulin localized at the peripheral shell of these neurites, with fewer integrations in the microtubule of the central core (Movie S2). Interestingly, while many putative growing plus-ends were detected among the parallel microtubules in the neurite shell, the rarer ones seen in the core often belongs to more curved microtubules, with relatively more integrated tubulin dots belonging to in-lattice insertion (Movie S2). We also observed more tubulin integration in the shell of neurites in more mature stage 2a neurons, with the tubulin integration at the neurite shell being frequently asymmetric with dorsal or lateral accumulations, and less tubulin integration at the ventral contact with the substrate (Fig. 6). Altogether, assessment of microtubule turnover and localization of newly integrated tubulin sites demonstrates that in-lattice tubulin incorporations occur in neuronal microtubules, including acetylated ones. This is clear in thicker neurites, thanks to the segregation between the core of acetylated microtubules and peripheral non-acetylated microtubules.

**Figure 6.**
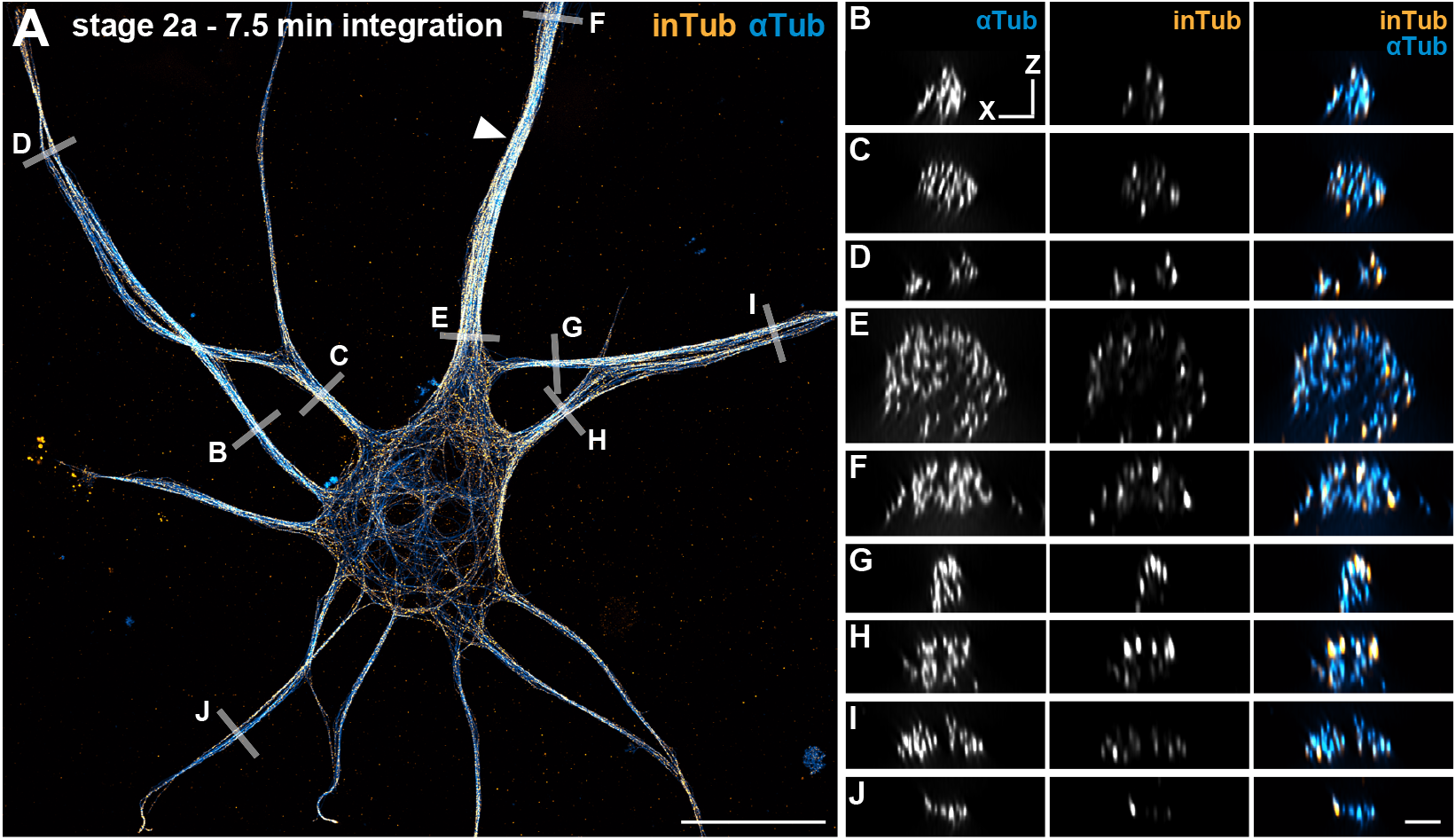
Microtubules show more turnover at the peripheral shell of neurites, and less within the neurite core. **A**. Deconvolved super-resolution spinning disk maximum intensity projections of a 1 DIV neuron at stage 2a, microinjected and fixed after 7.5 minutes of tubulin integration, processed for U-ExM and stained for α-tubulin (αTub, blue on overlays) and integrated tubulin (intTub, orange on overlays). Solid white arrowhead identifies the longest neurite. Scale bars, 10 µm after correction for the expansion factor. **B-E**. XZ sections across the neurites along the segments indicated in A showing the radial pattern of tubulin integration within thin, thick, proximal and distal neurites (D and I are section across two adjacent thin neurites). Scale bars, 0.5 µm.

## Discussion

In this work, we aimed at directly assessing microtubule turnover in developing hippocampal neurons by visualizing tubulin integration with super-resolution fluorescence microscopy shortly after microinjection of labeled tubulin. Microinjection of labeled tubulin in neurons has been achieved in previous works to evidence the microtubule transportation and polymerization in axons (Lim et al., 1990; Okabe and Hirokawa, 1990; Li and Black, 1996; Wang et al., 1996; Yu et al., 1996; Dent et al., 1999). Here, we focused on short time frames (∼10 minutes) and performed high-resolution imaging to reveal the fast in-lattice integration of tubulin as part of the turnover mechanisms of microtubules, taking advantage of the defined developmental stages of cultured rat hippocampal neurons (Dotti et al., 1988). In our microinjected neurons, the neurite median length was generally lower than 100 µm, a distance inferior to the 105 µm previously found reached by microinjected tubulin in 3 min along axons (Li and Black, 1996), likely alleviating the concern of bias from the diffusion time of microinjected tubulin in our results. This is strengthened by our consistent detection of tubulin integration at the tip of growing neurites (see Fig. 4A). At the whole-neuron level, we found that a significant amount of injected tubulin was integrated in the microtubule network within minutes, reaching a plateau after about 10 minutes. This suggests that developing neurons have a rather dynamic network, with the microtubule stabilization usually associated with neurons occurring at later stages (Baas et al., 2016).

We previously used microinjection of tubulin in mammalian non-neuronal cells and, by using Single Molecule Localization Microscopy (SMLM), demonstrated that in-lattice incorporation of tubulin occurs in parallel with active polymerization at plus-ends (Gazzola et al., 2023). SMLM was suitable because the microtubule network of these cells is sparse enough to visualize individual microtubules over several microns. In contrast, the microtubule arrays found within neurons are tightly bundled, leading to poor penetration and restricted antibody labelling. Therefore in this study, we implemented ExM in combination with super-resolution confocal microscopy to improve both resolution and labelling efficiency through uncrowding of these dense structures. The gain in resolution offered by U-ExM allowed us to unambiguously visualize instances of tubulin turnover, but we could not assess it over long distances on individual microtubules, which would be required for a quantitative assessment of in-lattice integration vs end renewal in distinct compartments. Also, ExM by nature is prone to introducing breaks in microtubules, potentially impeding the localization of true ends or microtubule nucleation sites along neurites. Furthermore, we are injecting small volumes, resulting in a low amount of injected tubulin relative to the endogenous tubulin pool (∼10% vs 90%, Okabe and Hirokawa, 1988). This most likely explains the dispersed labelling of integrated injected tubulin at high resolution and the dotted appearance of plus-ends in ExM images (see Fig. 5,6 and Movie S2). It is also possible that the integration of the endogenous tubulin pool is faster than that of exogenous microinjected tubulin owing to tubulin modification. However, with the rapid diffusion of injected tubulin across these small neurons, we are still likely to detect integration events with reasonable sensitivity. Indeed on ExM images we could identify both stretches of microinjected tubulin as well as sparse well dispersed spots likely corresponding to plus-end and in-lattice incorporations respectively. This presence of in-lattice integration adds a layer of complexity to the already ambiguous notion of microtubule stability, as it dissociates it from the lifetime of its tubulin components: akin to the ship of Theseus, a long-lived microtubule can be one that is subjected to a high rate of in-lattice turnover (Guedes-Dias and Holzbaur, 2019).

The fact that we were able to observe early in-lattice integrations not only in non-acetylated microtubules but also in acetylated microtubules indicates that these integrations may contribute to what characterizes an acetylated microtubule as being long-lived and stable (Janke and Magiera, 2020; Lu, 2025). We observed tubulin turnover to be sparser in acetylated microtubule bundles, in line with the idea that accumulation of post-translational modification requires stability of tubulin dimers themselves in the lattice. This difference between non-acetylated and acetylated microtubules was easier to detect in thick neurites, where they segregate between a peripheral shell and a denser core. We found a high density of integration sites within the peripheral shell, while they were more sparsely distributed in the core bundles. This is consistent with the distinction between two microtubule subsets in dendrites based on posttranslational modifications of α-tubulin, with tyrosinated dynamic and acetylated non-dynamic microtubules located at the shell and core of dendrites, respectively (Katrukha et al., 2021; Iwanski et al., 2025). The distinct patterns of tubulin integrations we observe in neurites are direct evidence that, indeed, microtubules at the shell are highly dynamic via active incorporation of tubulin at their plus-ends in addition to in-lattice repair, and the more stable microtubules at the core have sporadic turnover of their lattice.

An important result is that the longest neurite, be it at asymmetrical stage 2 or at stage 3 when we can assume the axon is specified, showed a reduction in the rate of tubulin turnover, while the minor neurites had higher turnover (see Fig. 4). This finding is consistent with the local increase in microtubule acetylation in the developing axon compared to other neurites (Shea, 1999; Witte et al., 2008). It directly demonstrates that the polarization of neurons depends on or is concomitant with the appearance of a locally stabilized population between the future axon and dendrites. Increase of neuronal microtubule stability by exposure to paclitaxel induces the formation of multiple axons, which corroborates a microtubule stability-dependent mechanism of neuron polarization (Witte et al., 2008). Of note, microtubule stabilization influences the dynamics of the nearby actin cytoskeleton, so that it might act indirectly via actin network re-organization (Bradke and Dotti, 1999).

By directly assessing the turnover of neuronal microtubules, our study demonstrates the existence of in-lattice integration in addition to plus-end dynamics. It further grounds the notion that microtubules, via subtle changes in their stability, participate in the specification of the axon and the emergence of neuronal polarity. This contributes to building an evolving model where the intrinsic renewal and specific conformational changes of the microtubule lattice, and their cross-talks with MAPs and motors (Shima et al., 2018; Chew and Cross, 2025), drive neuronal polarization and stabilize the nascent axon.

## Acknowledgments

We would like to thank Laurent Blanchoin (CytoMorpho lab, Grenoble) and Tamiko Ishizu for discussions, as well as Marie-Jeanne Papandréou, Louis Romette and the NeuroCyto lab members. We thank Erik W. Dent (University of Wisconsin-Madison) for help on the micro-injection approach; Bernard La Scola (Mephi, IHU, Marseille) and Jean-Pierre Baudoin (electron microscopy facility, IHU, Marseille) for help with the SEM images of capillaries; Bertrand Vernay (NCIS imaging facility, INP, Marseille) for help with image acquisition.

The project leading to this publication has received funding from the French National Research Agency (ANR-24CE13-7996 “MicRON” to C.L and M.T.), from the French government under the “France 2030” investment plan managed by the French National Research Agency (NeuroSchool ANR-16-CONV000X / ANR-17-EURE-0029), and from Excellence Initiative of Aix Marseille University - A*MIDEX (DINoMIR AMIDEX AMX-22-RE-AB-137 to C.L., Neuro-Marseille AMX-19-IET-004). We would like to thank the Neuro-Cellular Imaging Service and Nikon Center for Neuro-NanoImaging at INP, with funding from CPER-FEDER (Plateforme NeuroTimone PA0014842) and the Institut Marseille Imaging (AMX-19-IET-002) for complementary equipment funding from Excellence Initiative of Aix-Marseille University – A*MIDEX, a French “Investissements d’Avenir” program (ANR-11-IDEX-0001).

## Author Contributions

**Ciarán Butler-Hallissey:** Conceptualization, Methodology, Investigation, Analysis, Data Curation, Writing - Original Draft, Writing - Review and Editing, Visualisation. **Harrison York:** Analysis, Writing - Review and Editing, Visualisation. **Florence Pelletier:** Methodology. **Jean-Marc Goaillard:** Methodology, Analysis, Resources, Writing-Review and Editing. **Jérémie Gaillard:** Resources. **Manuel Théry:** Conceptualization, Resources, Writing - Review and Editing, Supervision and project administration, Funding acquisition. **Pascal Verdier-Pinard:** Analysis, Writing - Original Draft, Writing - Review and Editing, Visualisation. **Christophe Leterrier:** Conceptualization, Analysis, Writing - Original Draft, Writing - Review and Editing, Visualisation, Supervision and project administration, Funding acquisition.

## Methods

### Sample preparation and imaging

#### Animals and neuronal cultures

The use of Wistar rats followed the guidelines established by the European Animal Care and Use Committee (86/609/CEE) and was approved by the local ethics committee (agreement D13-055-8). Rat hippocampal neurons were cultured following the Banker method, above a feeder glia layer (Kaech & Banker, 2006). Hippocampi from E18 rat pups were dissected and homogenized by trypsin treatment followed by mechanical trituration. Cells were counted and seeded on the poly-L-lysine-coated (Sigma #P2636) coverslips (18 mm-diameter round #1.5H, Marienfeld #0117580) at a density of 4,000-8,000 cells/cm^2^ for 3 h in serum-containing plating medium. Coverslips were then transferred, cells down, to petri dishes containing confluent glia cultures conditioned in B27-supplemented (Thermo Fisher #17504044) neurobasal medium (NB^+^, Thermo Fisher #21103049). Neurons were microinjected and fixed at 1 day in vitro (1 DIV), between 22-27 h after plating.

#### Neuron developmental stage definition

Neuronal development stages were defined according to the established stages (Dotti et al., 1988) with a splitting of stage 2 between stage symmetric (stage 2s) and stage 2 asymmetric (stage 2a). Stage 2 symmetric neurons displayed neurites with similar length, while stage 2 asymmetric neurons would display an elongated neurite that was 1.5-2 times longer than other neurites, but still showed a less defined neurite maturity compared to stage 3 neurons, where there were short neurites with one neurite longer than 50 μm and 2-3 times longer than other neurites (Fig. S3A).

#### Tubulin purification and labelling

Biotinylated tubulin was prepared as previously described (Shelanski, 1973; Hyman et al., 1991; Vantard et al., 1994). Briefly, tubulin was purified from fresh bovine brain by three cycles of temperature-dependent assembly and disassembly in Brinkley buffer 80 (BRB80 buffer: 80 mM PIPES pH 6.8, 1 mM EGTA and 1 mM MgCl_2_, 1 mM GTP). MAP-free brain tubulin was purified by cation-exchange chromatography (EMDSO, 650M, Merck) in 50 mM PIPES, pH 6.8, supplemented with 1 mM MgCl2 and 1 mM EGTA. Purified tubulin was obtained after a cycle of polymerization and depolymerization. Finally, tubulin was labelled with EZ-Link sulfo-NHS-LC-LC-biotin, resuspended in microinjection buffer (50 mM potassium glutamate, 1 mM MgCl2, pH 6.8), aliquoted in 500 µL plastic tubes, flash frozen in liquid nitrogen and stored at -80°C.

#### Microinjection of neurons

Microinjection capillaries with an outer diameter of 234 ± 19 nm (Fig. S1A) were pulled from thick-walled borosilicate glass with filament (GC150F-10, Harvard Apparatus) on a DMZ-Universal Puller (Zeitz Instruments). Biotinylated tubulin stored at -80°C was diluted to a working solution of 30 µM by diluting it in a 0.22 µm filtered microinjection solution containing 0.7 µM TRITC-dextran MW 70K (Thermo Fisher, #D1819), 50 mM potassium L-glutamate, 0.5 mM magnesium sulphate, pH 6.8. Capillaries were positioned needle end down at an acute angle on adhesive putty. A small volume of microinjection solution was added to the top end of the capillary, filling approximately 3 mm of the capillary using a microloader tip (Calibre Scientific). Due to capillary forces, the solution travelled down to the needle end. After 2 min, additional microinjection solution was added by inserting the microloader into the capillary, filling it to about 1 cm from the needle end.

Neuron coverslips at 1 DIV were removed from the culture dish, paraffin dots were removed, and the coverslip was mounted in a custom-made imaging chamber. The chamber was then filled with 2 mL NB^+^. On the external surface of the coverslip, a lipid PAP pen was used to mark a ring within which the neurons would be injected.

Microinjection was performed on an InjectMan 4 micromanipulator (Calibre Scientific). Before injecting neurons, the needle was lowered into a dish and allowed to flow for 5 min at 200 hPa to ensure sufficient flow of tubulin. Neurons were injected using a continuous pressure of 30 hPa. When required, pressure was adjusted to ensure an optimal low flow rate. Injections were performed on a Zeiss Axio Observer.Z1 with a 63x DIC air objective NA 0.75. Successful microinjection was apparent when FITC-dextran could be seen entering the neuron immediately after this, the needle was removed, and cells were viewed with DIC to verify that viability was maintained (Movie S1).

Tubulin integration time in injected neurons was determined by synchronizing a timer with a video recording of the experiment (Fig. S1B). At the end of the experiment, neurons were permeabilized with 0.25% Triton-X-100 (Millipore Sigma, #T8787) and 4% sucrose in PEM at 37°C for 15 seconds, then fixed with the permeabilization buffer supplemented with 3% PFA (Delta Microscopy, #EM-15714), 0.05% glutaraldehyde (Millipore Sigma, #3G5882) for 10 min. Coverslips were then washed three times with phosphate buffer.

#### Fluorescence immunocytochemistry

Neurons were blocked for 2 h with 0.22% gelatine (Millipore Sigma, A7030), 0.1% Triton X-100 in phosphate buffer. Using the same blocking buffer for antibodies and washes, neurons were incubated with a goat anti-biotin antibody (Thermo Fisher, #31852) for 1 h. After 3 washes, donkey anti-goat Alexa Fluor 647 Plus (Thermo Fisher, #A32849) was then incubated for 1 h. After 3 washes, neurons were incubated with rabbit anti α-tubulin (Abcam, #Ab18251) and mouse anti acetylated α-tubulin (Millipore Sigma, #T7451) antibodies overnight at 4°C. The next day, samples were washed 3 times and incubated with donkey anti-rabbit Alexa Fluor 555 (Thermo Fisher, #A-31572) and donkey anti-mouse Alexa Fluor 488 (Thermo Fisher, #A21202) antibodies. Following three washes with phosphate buffer, coverslips were mounted onto slides with Prolong glass mounting medium (Thermo Fisher #P36980).

#### Expansion Microscopy

Following a modified version of U-ExM (Gambarotto et al., 2019), fixed neurons were incubated with samples in 1.4% PFA and 2% acrylamide (Thermo Fisher #J62480.AP) in 1x PBS for a minimum of 1 h at 37°C, then allowed to cool before being put on ice. A parafilm bottom chamber was prepared, and an adhesive silicone sheet (Grace Biolabs #GBL665581) with a 4 mm diameter well was firmly attached.

When embedding each coverslip, a 90 µL aliquot of frozen monomer solution of 19% sodium acrylate (Millipore Sigma, #408220), 10% acrylamide, 0.1% N,N′-methylenebisacrylamide (Millipore Sigma, #M1533) in 10x PBS was thawed with 5 µL 10% TEMED (Millipore Sigma, #T9281) and 5 µL APS (Millipore Sigma, #A3678) added and mixed. Rapidly, 40 µL of the solution was added into the 4 mm well, and the coverslip was mounted neuron side down over the well, centering the PAP pen ring inside the boundaries of the 4-mm well. Once mounted, coverslips were kept on ice for 5 min before being placed in an incubator for 1 h at 37°C.

The polymerized gels and coverslip were then removed from the silicon well and placed in denaturing buffer (200 mM SDS, 200 mM sodium chloride, 50 mM Tris Base in MilliQ water, pH 9) for 30 min to detach the gels from the coverslip. The detached gels were then placed in fresh denaturation buffer and denatured in a heat block at 95°C for 1.5 h. Gels were then rinsed three times in 1x PBS. Immunostaining was performed in the same sequential order as non-expanded neurons, with primary antibodies incubated overnight on a rocker at 70 rpm and secondary antibodies for 2 h with 30 min wash steps. After immunostaining, the gel was fully expanded by at least three rounds of MilliQ water. Before imaging, gels were mounted on poly-L-lysine-coated glass-bottom dishes.

#### Super-resolution confocal imaging

All imaging was performed on a Nikon Eclipse Ti2-E inverted microscope with a Yokogawa CSU-W1 spinning disk confocal and SoRa super-resolution module (4x zoom lens). Laser and emission filters used were 641 nm, filter: 708/75nm (Alexa Fluor 647/Abberior Star Red), laser: 561 nm, filter: 600/52 nm (Alexa Fluor 555) and laser: 488 nm, filter: 525/30 nm (Alexa Fluor 488), captured with a Hamamatsu Fusion BT sCMOS camera.

Non-expanded neurons imaging was performed with CFI Apo TIRF 60XC oil-immersion objective, NA 1.49. For expanded neurons, imaging was performed with a C-Apochromat 63x water-immersion objective, NA 1.2.

Z stacks captured the entire neuron volumes using a Mad City Labs Nano-Z Piezo stage. Channels were captured sequentially, followed by a Z step of 150 nm (non-expanded neurons) or 180 nm (expanded neurons) for the following plane.

Qualitative images were processed using 3D deconvolution using NIS-Elements Advanced Research with 3D Deconvolution Richardson Lucy with a maximum of 10 iterations. To capture a whole neuron extending beyond a single field of view, additional fields were taken to capture the entire neuron, and later the tiles were stitched as part of a processing pipeline.

### Image processing and analysis

#### Tile stitching and pre-processing

Representative images were deconvolved and stitched with linear blending for the overlapping regions. Images were corrected for channel misalignments using a custom macro using multi-spectral beads as a reference (Laine et al., 2019; Pylvänäinen et al., 2023). The neurons captured with tiled imaging were then grouped and stitched using a custom macro that implemented pairwise stitching (Preibisch et al., 2009) using stage coordinates metadata (Fig. S2A). Images used for quantification were processed similarly, without the deconvolution step.

Machine-learning software ilastik (Berg et al., 2019), was used for 3D segmentation, with contrast increased to identify and include low-intensity acetylated tubulin in the segmentation training (Fig. S2A). Following segmentation of the images, all further processing and analysis were performed in Fiji (Schindelin et al., 2012). As the α-tubulin antibody displayed a weakened microtubule staining in the cell body, it was combined with the acetylated tubulin mask to create the total α-tubulin segmentation mask (Fig. S2B). To create compartment ROIs, maximum intensity projections were created, and a custom macro was used to create an outline of each neuron that was manually split into compartments of cell body, neurites and lamellipodia before saving them as ROIs (Fig. S2C).

#### Integration rate measurements for total and acetylated tubulin

To assess the rate of integration of microinjected tubulin into the neuron (Fig. 1, 2 and S4), we used the binary segmentation masks of total tubulin (totTub) generated by ilastik. This allowed us to determine whether a pixel was positive for α-tubulin (αTub), acetylated α-tubulin (acTub), or integrated tubulin (intTub), ignoring differences in signal strength between two pixels that were both positive. We then calculated the fraction of integrated tubulin-positive pixels overlapping with total tubulin or only acetylated α-tubulin pixels at the whole neuron level. To identify these overlapping pixels between channels, the segmented binary images of integrated tubulin were processed against total tubulin and acetylated tubulin using the AND function in Fiji’s image calculator. The integration rates were determined dividing the number of overlapping pixels by the number of pixels positive for total tubulin or acetylated α-tubulin (output 1, Fig. S2B).

Intensity distribution measurements. For intensity-based measurements (Fig. 3, 4 and S5), the total tubulin segmentation mask was used to identify all pixels of interest. Pixels outside the total tubulin mask were converted to 0 for all channels. A sum intensity projection was then performed, and the total signal was measured for each channel across the neuron compartments (Fig. S2C). The raw integrated density was calculated within each compartment ROI, and we then calculated the “distribution percentage” for this label, expressed as a percentage of the total integrated density for the set of ROIs representing the entire neuron. These distribution percentages provided normalized measurement of the distribution of each label in each compartment, facilitating comparisons of relative signal distribution patterns between neurons (output 2, Fig. S2E). From these distribution percentages, we could then calculate the “distribution ratio” for acetylated α-tubulin and integrated tubulin that measures the relative enrichment of acetylation and integration compared to the quantity of microtubules in each compartment. These distribution ratios were obtained by dividing the distribution percentages for acetylated α-tubulin and integrated tubulin by the distribution percentage of α-tubulin (outputs 3 and 4, Fig. S2E).

#### Neurite length measurement

Neurite lengths (see Fig. S5) were measured using a 2D geodesic distance map via the chess-knight method from the Fiji plugin MorphoLibJ (Legland et al., 2016). A binary image of the cell body ROI served as the “marker”, while a binary image of all non-cell body ROIs functioned as the “mask.” To convert the distance map into scaled distances, the pixel values were multiplied by the pixel size (0.028 µm for 4x SoRa images). The maximum distance within each ROI was then measured to determine the maximum neurite length (output 5, Fig. S2F).

#### Movie preparation

Movie S2 was rendered volumetrically in Imaris 10.2. To compare the spatial pattern of tubulin integration the outer 300 nm of the neurite was segmented in 3D using MorpholibJ (Legland et al., 2016) to distinguish the peripheral shell and the remaining central core.

#### Quantification analysis and statistics

Intensity measurements and associated data files were transformed and processed in Microsoft Excel, using Power Query and pivot tables for data organization. Finalized tables were plotted and statistically analyzed using GraphPad Prism. To assess data distribution, the Shapiro-Wilk test was applied. For comparisons between two groups, an unpaired t-test was used when data followed a normal distribution, while the Mann-Whitney U test was used for non-normally distributed data. For comparisons involving three or more groups, Welch’s ANOVA was applied for normally distributed data with unequal variances, followed by Dunnett’s T3 multiple comparisons post-hoc test. For non-normally distributed data, the Kruskal-Wallis test was used, followed by Dunn’s multiple comparisons post-hoc test. Differences between Padé Approximant (1,1) fits for integration of total and acetylated tubulin were assessed using the extra sum-of-squares F-test (α=0.05). Significances are reported on the graphs as follow except when specified: ns non-significant; * p<0.05; ** p<0.01; *** p<0.001.

## Supplementary material

**Figure S1.**
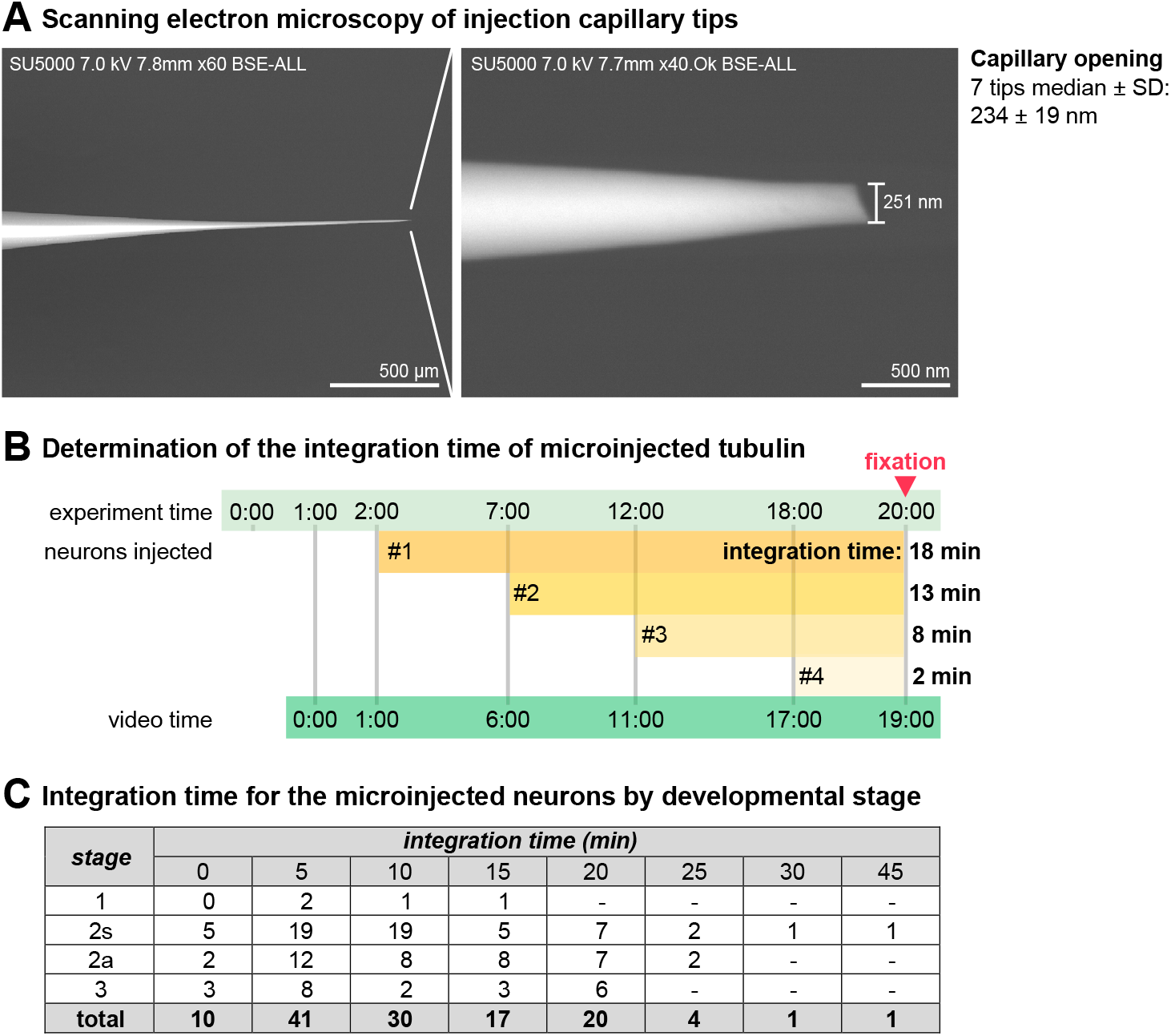
Microinjection pipettes and timing of tubulin integration after microinjection. **A**. Scanning electron microscopy image of a microinjection capillary tip showing an opening of 251 nm, and statistics of the opening over 7 pipettes. **B**. Methods to determine the integration time of microinjected tubulin. The microinjection procedure is continuously recorded (screen capture), allowing to precisely measure the time between the injection of each neuron and the fixation, as well as identify microinjected neurons after fixation and labeling. **C**. Summary table of the integration time intervals for each of the 124 microinjected neurons divided by developmental stage.

**Figure S2.**
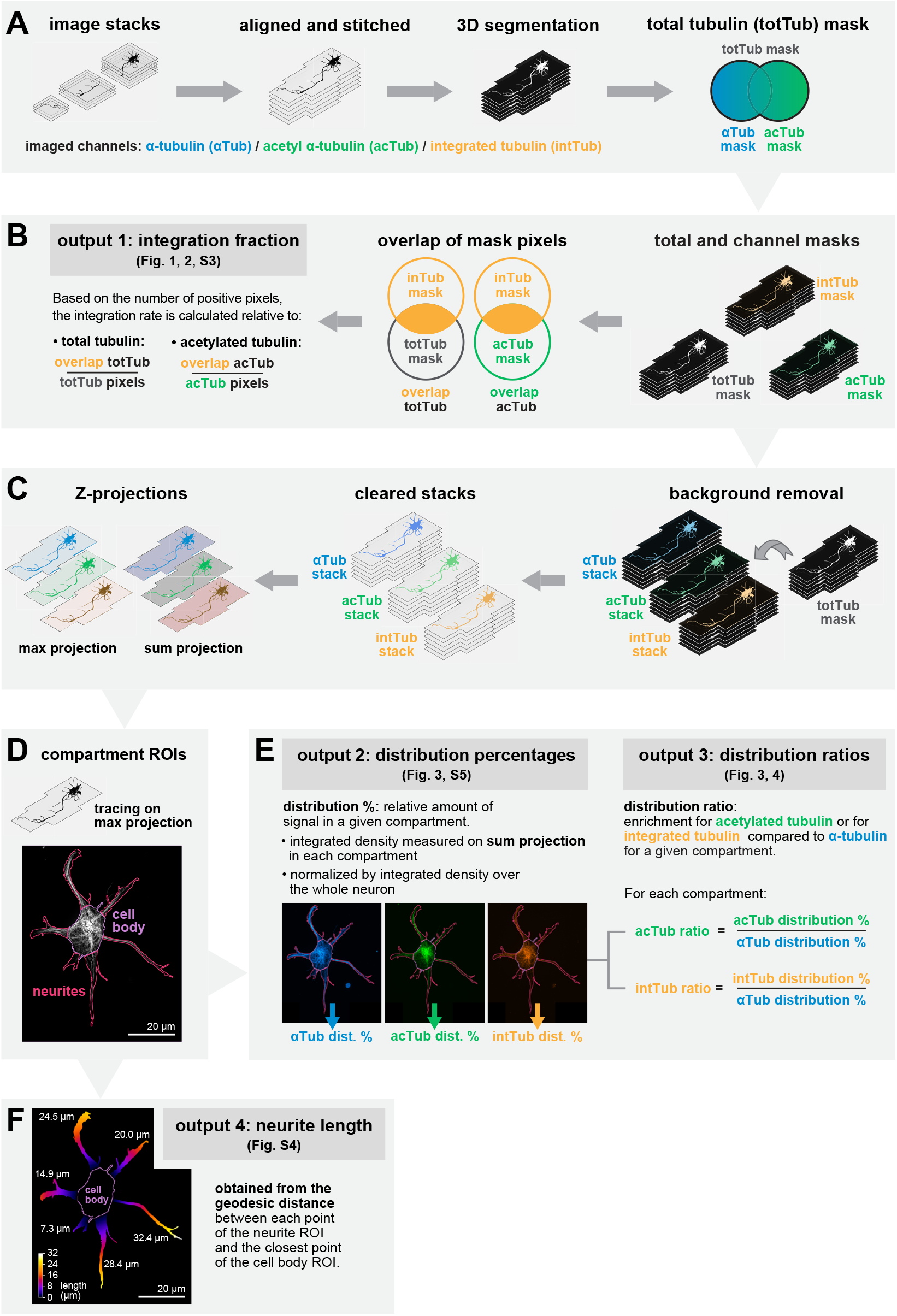
Image processing and analysis workflow. **A**. Whole neurons were captured as several manually overlapping image stacks using a 60X objective (left), and the stacks were then corrected for chromatic aberration, aligned and stitched (center left). Each channel of the stitched images was segmented in 3D to generate segmentation masks (set of pixels positive for each channel, center right), then a total tubulin (totTub) mask was created by combining (OR operation) the segmentation masks of α-tubulin (αTub) and acetylated α-tubulin (acTub, right). **B**. From the segmentation masks (right), the injected tubulin (intTub) positive pixels that were also positive for total tubulin (totTub) or for acetylated α-tubulin (acTub) were determined (AND operation, center). The number of these overlapping pixels was then divided by the totTub or acTub pixels, respectively, to determine the integration fraction relative to totTub or acTub (output 1), as shown in the graphs of Fig.1 and 2 (left). **C**. For intensity-based measurements in each channel, the background was first removed by multiplying each channel by the totTub mask (right). This resulted in the background pixels being converted to zero while preserving the original intensity values of the foreground pixels (center). The background cleared stacks were then projected along the Z axis to create maximum and sum projections (left). **D**. The maximum projection was used to create the neuron outline, which was then manually divided into compartments (cell body, neurites, and lamellipodia). **E**. For each channel, the signal within each compartment was measured as the integrated density on the sum projection image and expressed as distribution percentages (output 2) after dividing by the integrated density over the whole neuron. The distribution percentage measures the relative amount of signal for a given channel in each compartment, as shown in Fig. 3 and S5. Next, the distribution ratios (output 3) for acetylated α-tubulin (acTub) and integrated tubulin (intTub) were calculated for each compartment by dividing the corresponding distribution percentage by the distribution percentage of the α-tubulin (αTub). The distribution percentage thus measures the relative enrichment for acTub or for intTub in each compartment, as shown in Fig. 3 and 4. **F**. From the segmented neurites and cell body, the length of each neurite was determined by using the geodesic distance between ROIs (distance between each point of the neurite ROI and the closest point of the cell body ROI), considering the maximum geodesic distance value in μm as the neurite length.

**Figure S3.**
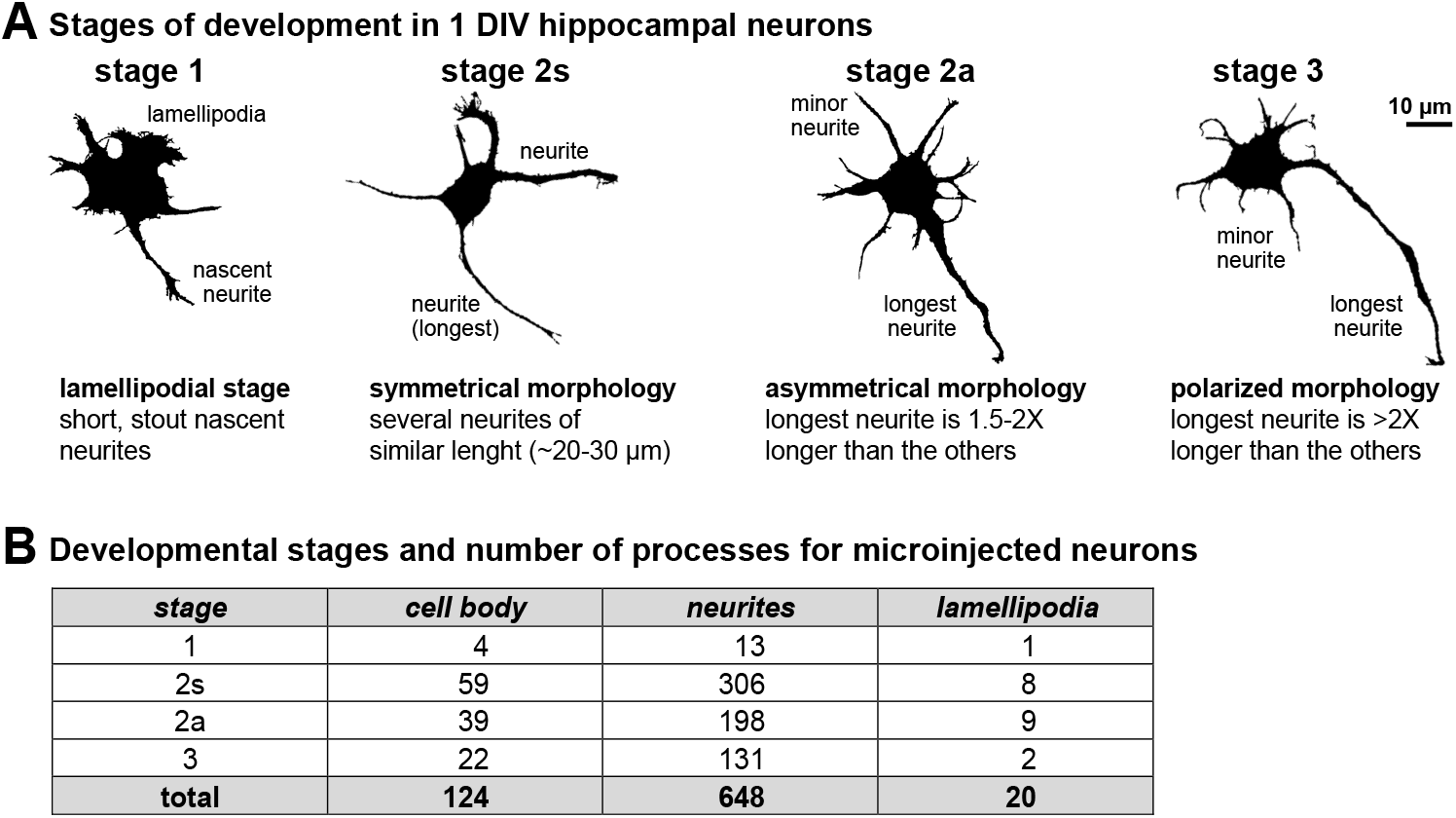
Developmental staging of hippocampal neurons. **A**. Silhouette of microinjected neurons illustrating the 4 stages of development present in 1 DIV cultures:1, 2s, 2a and 3, with criteria for stage decision. **B**. Summary table of the microinjected neurons developmental stage and compartments: cell body, neurites and lamellipodia. Due to the low occurrence of stage 1 neurons and lamellipodia, they were not considered further in the analysis.

**Figure S4.**
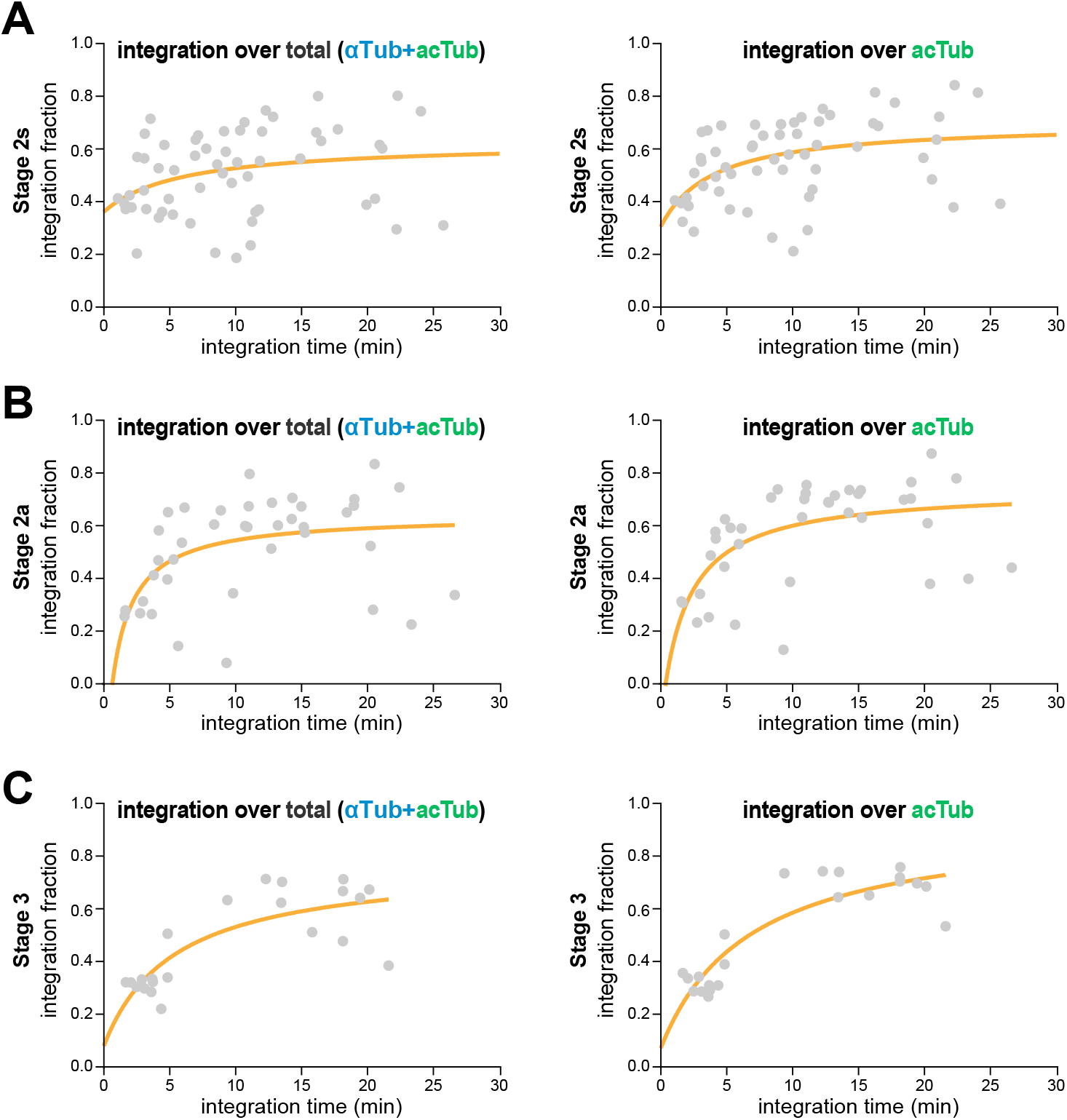
Integration fraction of microinjected tubulin at each developmental stage. **A**. Graph of the integration fraction of microinjected tubulin for the injected neurons at developmental stage 2s averaged over all integration times. The integration fraction as a function of time is calculated over the total tubulin positive pixels (left) or over the acetylated α-tubulin positive pixels (right). Orange curve is a fit using nonlinear regression (Padé (1,1) approximant equation). N = 59 neurons. **B**. Graph of the integration fraction of microinjected tubulin for the injected neurons at developmental stage 2a averaged over all integration times, similar to A. N = 39 neurons. **C**. Graph of the integration fraction of microinjected tubulin for the injected neurons at developmental stage 3 averaged over all integration times, similar to A. N = 22 neurons.

**Figure S5.**
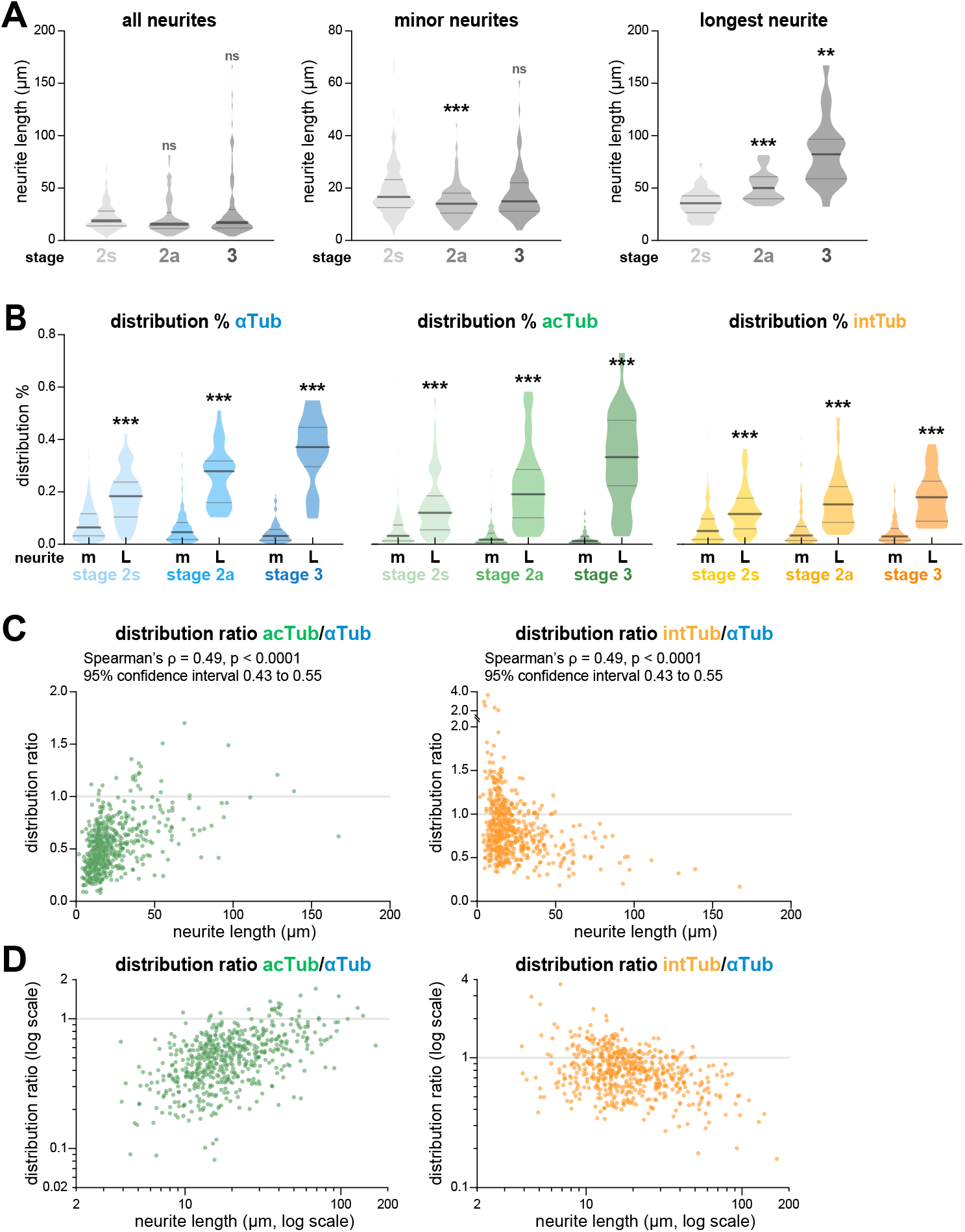
Growth of neurite length and relationship with tubulin acetylation and integration enrichment. **A**. Graphs of the lengths in µm for all neurites (left), minor neurites (center), and longest neurites (right) as a function of the developmental stage (2s, 2a and 3). Branched neurites were not included in the analyses. Violin plots show the median (thick horizontal segment) and upper 75th and lower 25th interquartile ranges (thin horizontal segments). Statistical analysis was performed using the Kruskal– Wallis nonparametric test followed by Dunn’s multiple comparisons test, with significance testing between a developmental stage and the one just to its left (previous stage). Stage 2s: all neurites N = 274, minor neurites N = 223, longest neurites N = 51; stage 2a: all neurites N = 171, minor neurites N = 136, longest neurites N = 35; stage 3: all neurites N = 116, minor neurites N = 96, longest neurites N = 20. **B**. Graphs of the distribution percentages for α-tubulin (αTub, left), acetylated α-tubulin (acTub, center) and integrated tubulin (intTub, right) in minor (m) and longest (L) neurites. Violin plots show the median (thick horizontal segment) and upper 75th and lower 25th interquartile ranges (thin horizontal segments). Statistical analysis was performed using the Kruskal–Wallis nonparametric test followed by Dunn’s multiple comparisons test, with significance testing between the longest neurites and the minor neurites at the same stage (just to its left). Stage 2s: minor neurites N = 223, longest neurites N = 51; stage 2a: minor neurites N = 136, longest neurites N = 35; stage 3: minor neurites N = 96, longest neurites N = 20. **C**. Scatter plots of the distribution ratios for acetylated α-tubulin (acTub, left) and integrated tubulin (intTub, right) for each neurite as a function of the neurite length. Spearman’s rank correlation coefficient (ρ) was calculated to assess the monotonic relationship between the two variables, with the results summarized on top of the plot. **D**. Log transformation of the scatter plot in C, assessing the relationship between enrichment in tubulin acetylation (left) or tubulin integration (right) with neurite length.

**Movie S1.**
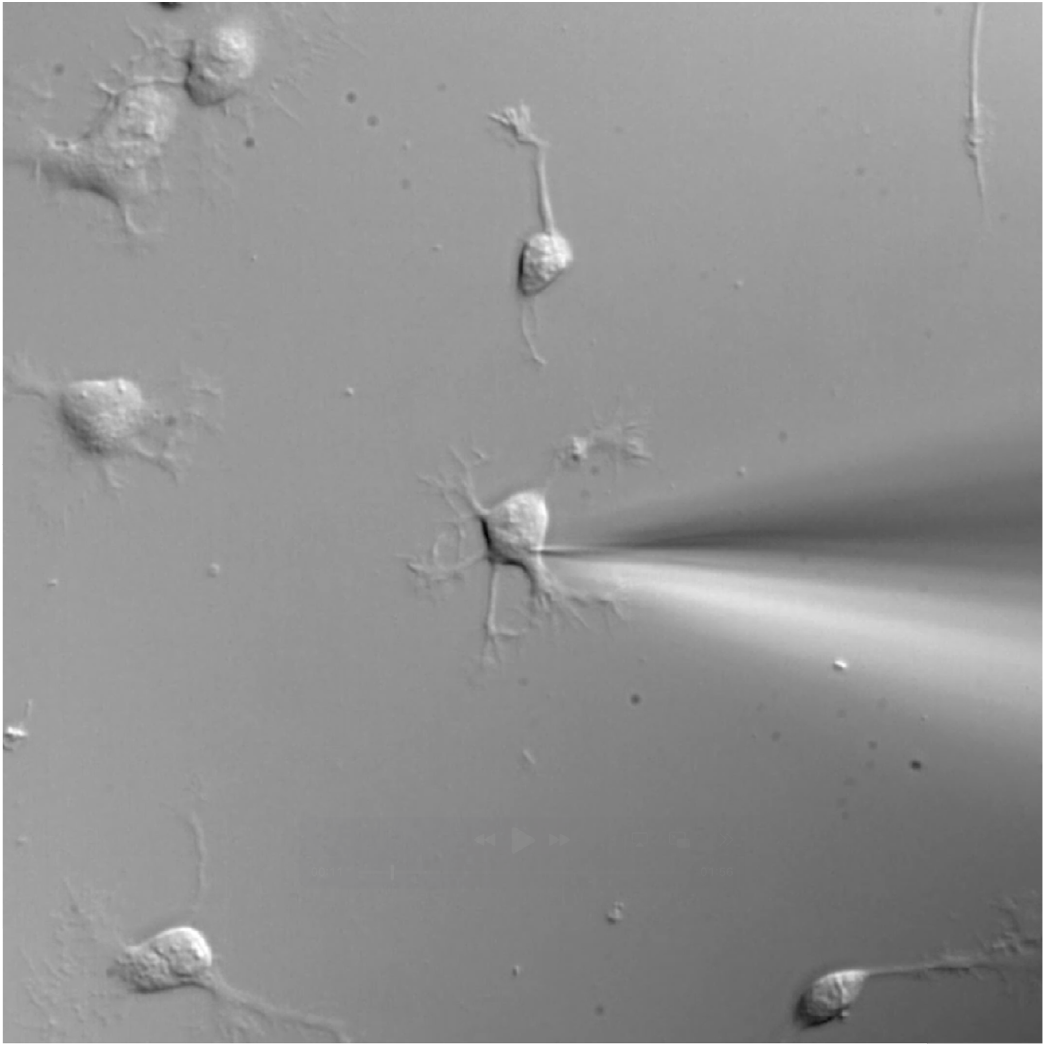
Live imaging of the microinjection procedure. Compilation of live DIC imaging showing the microinjection of 1 DIV rat hippocampal neurons with biotinylated tubulin and FITC-dextran, showing the successful microinjection and subsequent diffusion of FITC-dextran throughout the neuron.

**Movie S2.**
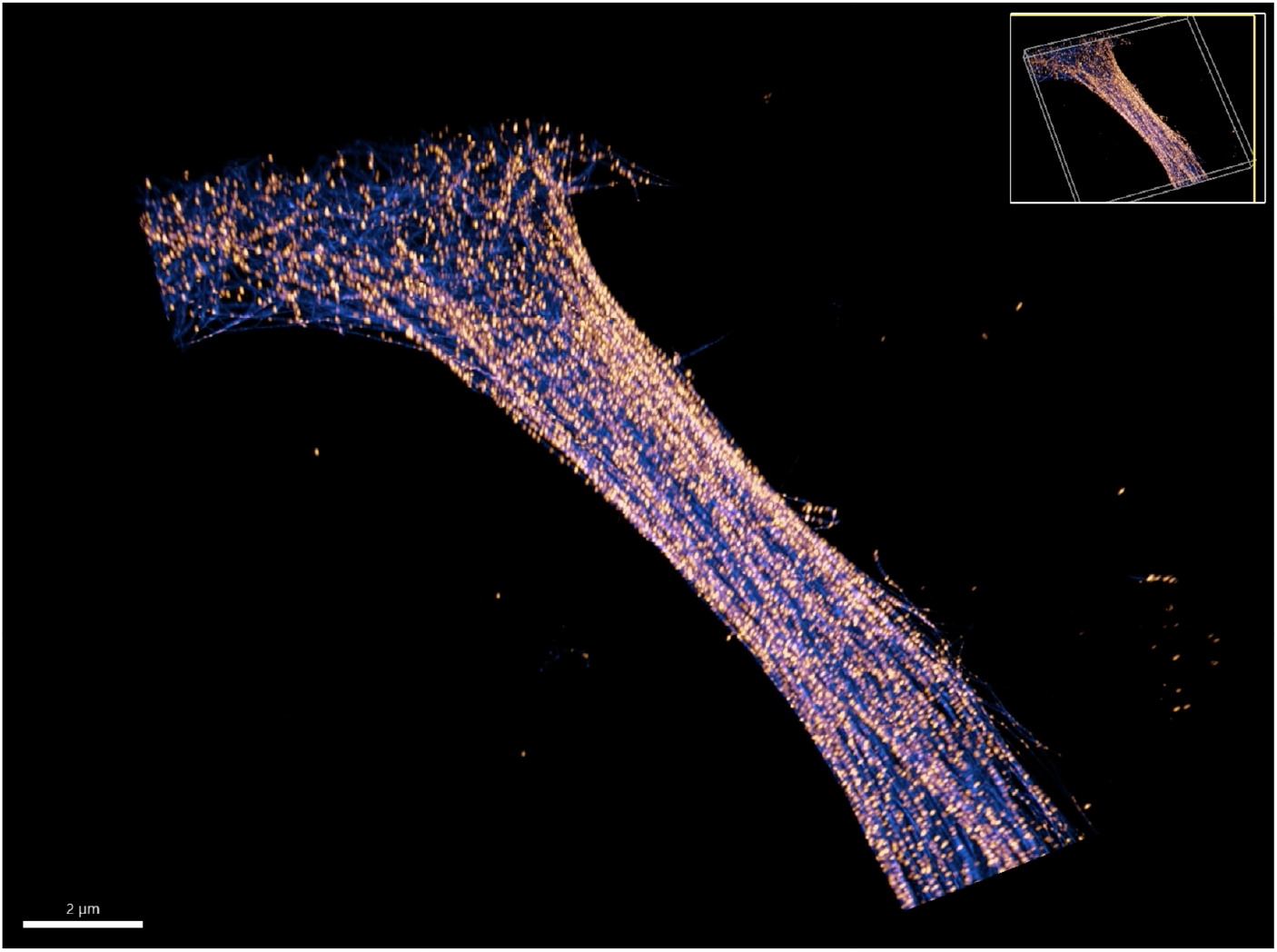
3D reconstruction of a microinjected, expanded neuron. Movie showing the 3D reconstruction from a super-resolution spinning acquisition of a 1 DIV neuron at stage 2s (same as in Fig. 5), microinjected and fixed after 5 minutes of tubulin integration, processed for U-ExM and stained for α-tubulin (αTub, blue) and integrated tubulin (intTub, orange). Annotations along the neurite highlight the presence of putative growing plus-end as strings of bright dots of integrated tubulin, and the presence of putative in-lattice integration events as fainter single dots along a microtubule. The second part of the movie splits the neurite into the peripheral shell and inner core domains, highlighting the parallel microtubules in the shell and the more curved morphology of the integration-rich microtubules in the core.

## Notes

### Competing Interest Statement

The authors have declared no competing interest.

